# Fast ontogenetic growth drives steep evolutionary scaling of metabolic rate

**DOI:** 10.1101/2021.03.29.437465

**Authors:** Tommy Norin

## Abstract

Metabolic rate (MR) changes with body mass (BM) as MR = *a*BM^*b*^, where *a* is a normalisation constant (log–log intercept) and *b* the scaling exponent (log–log slope). This scaling relationship is fundamental to biology and widely applied, yet a century of research has provided little consensus on why and how steeply metabolic rate scales with body mass. I here show that ontogenetic (within-individual) *b* can be strongly and positively related to growth rates of juvenile fish when food availability is naturally restricted, with fast growing individuals having steep and near-isometric metabolic scaling (*b* ≈ 1). I suggest that the steep evolutionary (among-species) scaling also found for fishes (*b* also approaching 1) is a by-product of natural selection for these fast growing individuals early in ontogeny, and that a weaker relationship between metabolic scaling and growth later in life causes variation in *b* at lower taxonomic levels (within orders or species). I support these ideas by showing that *b* within fish orders is linked to natural mortality rates of fish larvae.

## Introduction

Metabolic rate represents the cost of living and is related to many physiological and ecological processes (Schmidt-Nielsen 1984; Kooijman 2010; Sibly et al. 2012). Body mass profoundly affects metabolic rate so that larger animals generally have higher absolute metabolic rates than smaller animals. While this relationship sounds straightforward, how metabolic rate changes (scales) with body mass has been debated since at least 1932 (Kleiber 1932). There is general agreement that metabolic rate (MR) scales with body mass (BM) according to an allometric relationship: MR = *a*BM^*b*^, where *a* is a normalisation constant (the intercept on double-logarithmic axes) and *b* is the scaling exponent (the slope on double-logarithmic axes) (Rubner 1883; Kleiber 1932, 1947; Brody 1945; Schmidt-Nielsen 1984). This scaling relationship is fundamental to biology and widely applied, yet more than a century of research has failed to answer why metabolic rate scales with body size as it does, and why there appears to be variation in how steep this scaling is (the value of *b*) among different species or taxonomic groups. Knowing this is fundamentally important for understanding biological patterns and ecosystem dynamics, as it allows us to predict at what rates individuals and species of different sizes expend energy and need to consume resources. Metabolic scaling is also applied in influential size-based models predicting animal responses to climate change (Cheung et al. 2013; Deutsch et al. 2015, 2020), and in metabolic growth models predicting the maximum biomass that can be sustainably harvested from exploited populations (e.g. stock assessments in fisheries; von Bertalanffy 1957; Andersen 2019). The importance of understanding metabolic scaling is clear, yet it remains one of the most debated concepts in biology.

Metabolic rate was initially thought to scale in proportion to the surface area of animals, resulting in *b* ≈ ⅔ due to the need to dissipate internally-produced metabolic heat across the body surface (Rubner 1883), something that is still thought to apply to mammals and birds (White & Seymour 2003; White et al. 2006). After Rubner (1883), however, Kleiber (1932,1947) and Brody (1945) found an empirical value of *b* ≈ ¾ for mammals, which became largely fixed as a scaling law in biology (“Kleiber’s law”), backed by later fractal network theory explaining ¾-power scaling as arising universally from the need to distribute resources within the body through branching vessels (West et al. 1997, 1999; Banavar et al. 2010). While ¾-power scaling of metabolic rate is still widely promoted and applied (e.g. Brown et al. 2004, 2018; Sibly & Brown 2020), there is now increasing awareness that variation in *b* exists among different taxonomic groups (White et al. 2006), among endo- and ectotherms (White et al. 2007), across life’s evolutionary transitions (DeLong et al. 2010), and among different states of activity within the same animal (resting *vs*. active metabolic rates; White et al. 2007). There has also been a shift from explaining scaling as a purely physical and geometric phenomenon (West et al. 1997, 1999), to something that is being shaped by a suite of physiological, ecological, and evolutionary factors (Glazier, 2005, 2010, 2014a, 2018; Killen et al. 2010; Harrison 2017; Hatton et al. 2019). Overall, many researchers have embraced the variation that exists in *b* and are now trying to explain it for what it is, rather than explaining why it is not ¾.

Despite this reinvigorated interest in metabolic scaling, we still know surprisingly little about why there is variation in *b* among taxa and across different levels of biological organisation (Kozlowski & Wiener 1997; Kozlowski et al. 2003; Glazier 2005; DeLong et al. 2010; Hatton et al. 2019). This variation has received more attention in other branches of allometry (Gould 1966; Lande 1985; Cheverud 1982; Pélabon et al. 2013; Voje et al. 2013; Tsuboi et al. 2018) and is critical for understanding how evolutionary (among-species) patterns relate to patterns at ‘static’ (among individuals within species) and ontogenetic (among or within individuals at different life stages) levels, as ontogenetic scaling may constrain static scaling and both may constrain evolutionary scaling (Pélabon et al. 2013; Voje et al. 2013).

The evolution of metabolic allometry and its constraints have only recently been investigated extensively (White et al. 2019; Beaman et al., preprint), and ontogenetic scaling of metabolic rate has been investigated almost exclusively among different individuals at different life stages; variation in ontogenetic scaling within individuals – the source of natural selection – has received virtually no attention. Notable exceptions are a recent study on fish (Norin & Gamperl 2018) and an impressive phenotyping effort aimed at explaining heritable variation in the scaling of metabolic rate in insects (Beaman et al., preprint). The latter study found an absence of heritable genetic variation in the metabolic scaling exponent of cockroaches, and concluded that metabolic scaling can diversify among species, despite an evolutionary constraint within species (Beaman et al., preprint). However, studies investigating and comparing scaling patterns at different levels of biological organisation remain rare, but are needed to understand why there is variation in how metabolic rate changes with body mass.

Using published data on fish, I here investigate and compare how both standard (resting) metabolic rate (SMR) and aerobic maximum metabolic rate (MMR) scale with body mass among species (evolutionary scaling), among individuals within species (static scaling), and within individuals (ontogenetic scaling). I do this by combining phylogenetically-informed data on the scaling of SMR and MMR among 134 and 114 species of ray-finned fish, respectively, with the only existing data on within-individual scaling of metabolic rate in 68 and 33 individuals of two of these species (cunner, *Tautogolabrus adspersus*, and brown trout, *Salmo trutta*, respectively). I also examine if variation in growth rate among individuals is related to variation in metabolic scaling during ontogeny, and if ecological factors such as natural mortality rates may drive evolutionary scaling patterns.

## Methods

Data used here are available in the online supplementary material and analysed using R v. 4.0.2 (R Core Team 2020).

The dataset used to assess evolutionary (among-species) scaling of SMR and MMR is from Norin & Speers-Roesch (2020), which, in turn, is an updated version of the datasets from Killen et al. (2016, 2017). This dataset contains SMR and MMR values from 140 and 115 species, respectively, of juvenile or adult ray-finned fish (class Actinopterygii). Data on metabolic scaling of cunner (*Tautogolabrus adspersus*) are from Norin & Gamperl (2018), who measured the metabolic rates and body masses of 68 individuals in five separate trials over a 10-month period. Data for brown trout (*Salmo trutta*) are from Norin & Malte (2011), who measured the metabolic rates and body masses of 33 individuals in four separate trials over a 15-week period. All cunner and brown trout were kept and measured at the same temperature (15°C) in the original studies. The cunner were maintained on an *ad libitum* feeding regime and fed ~2.5% of their mean body mass daily (Norin & Gamperl 2018), while the trout were maintained on a restricted feeding regime and fed ~0.6% of their mean body mass daily (Norin & Malte 2011).

The data for brown trout were not originally analysed for scaling relationships and all data, including those for cunner, were therefore re-analysed here for consistency. I analysed ontogenetic (within-individual) and static (among-individual) scaling using linear mixed-effects (LME) models and within-subject centring (van de Pol & Wright 2009) in the packages *lme4* (Bates et al. 2015) and *lmerTest* (Kuznetsova et al. 2017). Each LME model had log_10_-tranformed whole-animal SMR or MMR as a response variable, log_10_-transformed body mass as predictor variable (fixed effect), and individual fish ID as a random effect. Body mass was included twice in each model to tease apart within- *vs*. among-individual relationships: each individual’s mean body mass across all five (cunner) or four (brown trout) trials was subtracted from its body mass at the time of each trial, and these mean-centred values were used as a fixed effect to express within-individual variation; the overall mean body mass of each individual (the same value repeated for each trial) was included as a separate fixed effect to express among-individual variation.

Evolutionary scaling relationships were analysed using phylogenetically-informed models. First, I extracted phylogenetic information from The Fish Tree of Life (Rabosky et al. 2018) and matched it to the species in my dataset using the package *fishtree* (Chang et al. 2019). There was no or dichotomous information available for six species, which were removed from the dataset, reducing it to 134 and 114 species for SMR and MMR, respectively. I then analysed these data using phylogenetic generalised least squares (PGLS) models in the package *ape* (Paradis & Schliep 2019). Each PGLS model had log_10_-tranformed whole-animal SMR or MMR as a response variable, while log_10_-transformed body mass and temperature at which metabolic rate was measured were included as fixed effects. Phylogenetic correlation was estimated using Pagel’s λ (Freckelton et al. 2002). Residuals from these PGLS models were used to adjust the SMR and MMR of each species to a common temperature of 15°C for graphical presentation.

Since metabolic rate is most often estimated as oxygen uptake rate, and researchers tend to favour smaller individuals or species that are more easily accommodated in respirometry chambers, it is possible that this potential bias towards smaller-sized animals could affect metabolic scaling relationships (Savage et al. 2004; Kolokotrones et al. 2010). To asses this, I extracted estimated asymptotic and maximum observed lengths for the species in the dataset from FishBase (Froese & Pauly 2019), and converted these to body masses using the length– weight relationships provided in FishBase. Since asymptotic and maximum observed sizes often differed, I took the mean of these two measures as a single measure of maximum body mass. I also calculated the relative body mass of each species in the dataset by dividing the body mass at which the species was measured for metabolic rate with its estimated maximum body mass. Maximum body mass estimates were only available for a subset of the species in the dataset (121 species for SMR, 109 for MMR), so I ran separate PGLS models on this reduced dataset to assess the influence of species-specific maximum and relative body mass on metabolic scaling. These PGLS models were structured as those above, except that either log_10_-transformed maximum body mass or log_10_-transformed relative body mass was included as an additional fixed effect.

Statistical differences between evolutionary, static, and ontogenetic mass-scaling exponents (*b*; model-estimated slopes) were evaluated at the p = 0.017 level to account for multiple comparisons (i.e. p = 0.05 divided by three comparisons within each of SMR and MMR). Pairwise comparisons between mass-scaling exponents for SMR (*b*_SMR_) and MMR (*b*_MMR_) within each of evolutionary, static, and ontogenetic levels were evaluated at the p = 0.05 level. Comparisons between static and ontogenetic mass-scaling exponents were done directly within each LME model, while all other comparisons were done by evaluating if the confidence interval for the difference between estimates of *b* did not contain zero.

The growth rates of cunner and brown trout presented here are specific growth rates from the original studies (Norin & Malte 2011; Norin & Gamperl 2018), and correlations (r) between metabolic rates and growth rates are Pearson’s.

To assess the possible influence of natural selection during early life stages on evolutionary scaling, I extracted estimates of natural mortality rates of fish larvae from the literature (McGurk 1986; Pepin 1991; Davis et al. 1991; Houde & Zastrow 1993). This was done for any species within teleost orders for which SMR or MMR data of at least six species were available in the metabolic rate dataset. Metabolic scaling exponents for these orders were estimated using PGLS models as above, except that only species from the relevant orders were included (99 species for SMR, 83 for MMR) along with an interaction between log_10_-transformed body mass and order. Mortality rates of species for which more than one estimate was available in the literature were averaged. Since very few mortality rate estimates existed for freshwater species, only data for marine species were included. Mortality rates of species from four orders (Gadidae, *Incertae sedis* in Eupercaria, Pleuronectiformes, and Scombriformes) matched the orders with six or more species in the metabolic rate dataset. Mortality rates also existed in the literature for species from the order Sparidae, which were also included in the analyses due to Sparidae’s close evolutionary proximity to *Incertae sedis* in Eupercaria (Fig. S1), despite metabolic rate data being available for only three species within Sparidae. The scaling exponents for SMR and MMR of Sparidae were assumed to be the same as those for *Incertae sedis* in Eupercaria, which seems reasonable (Fig. S1). Relationships between *b*_SMR_ or *b*_MMR_ and mortality rate at the order level were assessed using general linear regression.

## Results

Metabolic rate scales with body mass with exponents of 0.953 (95% CI: 0.894–1.012) for SMR and 0.878 (95% CI: 0.819–0.936) for MMR among the 134 and 114 species of ray-finned fish, respectively (evolutionary scaling; Fig. 1). The static (among-individual) scaling exponents for SMR are 0.869 (95% CI: 0.838–0.900) for cunner and 1.143 (95% CI: 0.886– 1.400) for brown trout; those for MMR are 0.941 (95% CI: 0.909–0.974) for cunner and 0.781 (95% CI: 0.603–0.959) for trout (Fig. 1). The mean ontogentic (within-individual) scaling exponents for SMR are 0.735 (95% CI: 0.706–0.764) for cunner and 0.271 (95% CI: 0.098– 0.438) for brown trout; those for MMR are 0.833 (95% CI: 0.811–0.856) for cunner and 0.693 (95% CI: 0.612–0.773) for trout (Fig. 1).

**Figure 1.**
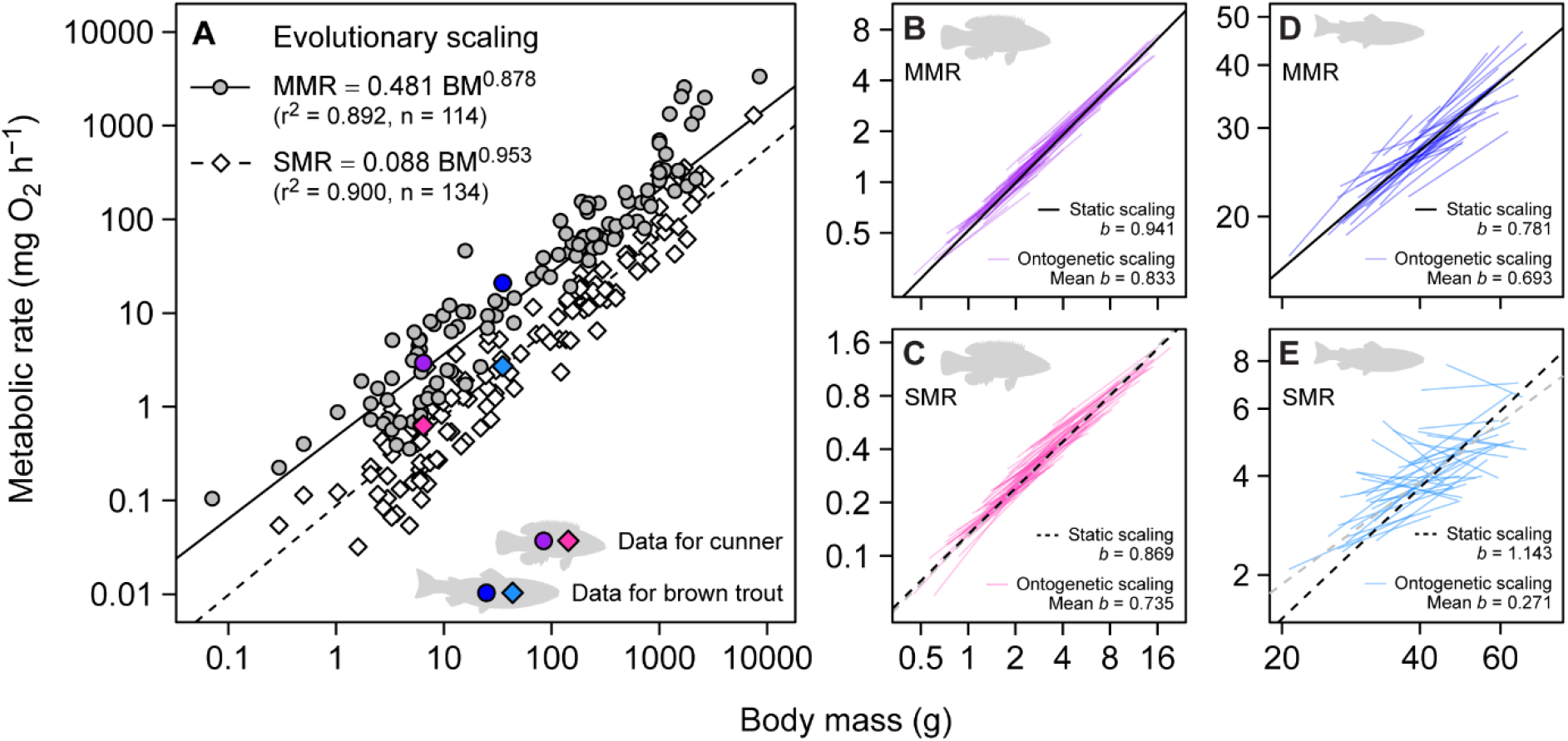
Scaling of metabolic rate with body mass at evolutionary, static, and ontogenetic levels in fishes. Evolutionary scaling (**A**) of metabolic rate with body mass (BM) is shown for standard metabolic rate (SMR) and maximum metabolic rate (MMR) of 134 and 114 species of ray-finned fish, respectively. Each data point represents a species, and the presented data have been adjusted to a temperature (T) of 15°C using residuals from the PGLS regression for SMR (log_10_ SMR = −1.386 + 0.953 log_10_ BM + 0.022 T; r^2^ = 0.893, λ = 0.869) or MMR (log_10_ MMR = −0.624 + 0.878 log_10_ BM + 0.020 T; r^2^ = 0.878, λ = 0.911). The coefficients for temperature translate to thermal sensitivities (*Q*_10_) of 1.66 for SMR and 1.60 for MMR. Scaling equations shown on the panel are for the temperature-adjusted data. Static and ontogenetic scaling (**B-E**) at 15°C are shown for the only two fish species for which comprehensive data exist on the scaling of metabolic rate within individuals (B-C: cunner, *Tautogolabrus adspersus*; D-E: brown trout, *Salmo trutta*). Mean scaling exponents (*b*) are noted on the panels and compared in Fig. 2, and full LME regressions are: (B) log_10_ MMR = −0.286 + 0.833 log_10_ BM_Ont_ + 0.941 log_10_ BM_Stat_ (_m_r^2^ = 0.968, _c_r^2^ = 0.977); (C) log_10_ SMR = −0.880 + 0.735 log_10_ BM_Ont_ + 0.869 log_10_ BM_Stat_ (_m_r^2^ = 0.949, _c_r^2^ = 0.956); (D) log_10_ MMR = 0.181 + 0.693 log_10_ BM_Ont_ + 0.781 log_10_ BM_Stat_ (_m_r^2^ = 0.726, _c_r^2^ = 0.797); and (E) log_10_ SMR = −1.260 + 0.271 log_10_ BM_Ont_ + 1.143 log_10_ BM_Stat_ (_m_r^2^ = 0.528, _c_r^2^ = 0.805). The coefficients for BM_Ont_ and BM_Stat_ represent the ontogenetic (within-individual) and static (among-individual) scaling exponents, respectively, and _m_r^2^ and _c_r^2^ are marginal and conditional r^2^, respectively. The mean static *b* for SMR of 0.89 recently reported for 16 fishes (Jerde et al. 2019) is included on panels C and E (grey dashed lines) for comparison with the static *b* for SMR of cunner and trout. Data are presented on logarithmic axes to linearise the relationship between metabolic rate and body mass.

Evolutionary scaling is always steeper (higher *b*) than ontogenetic scaling, although the difference is not significant for MMR of cunner (p = 0.159; Fig. 2). Static scaling is also steeper than ontogenetic scaling, except for MMR of trout (p = 0.381; Fig. 2). Evolutionary scaling is different from (steeper than) static scaling only for SMR of cunner (p = 0.014; Fig. 2). Scaling exponents for SMR and MMR are significantly different within both static and ontogenetic levels for both cunner and brown trout (p < 0.05), while *b*_SMR_ tends to be higher than *b*_MMR_ at the evolutionary level (p = 0.077) (Fig. 2).

**Figure 2.**
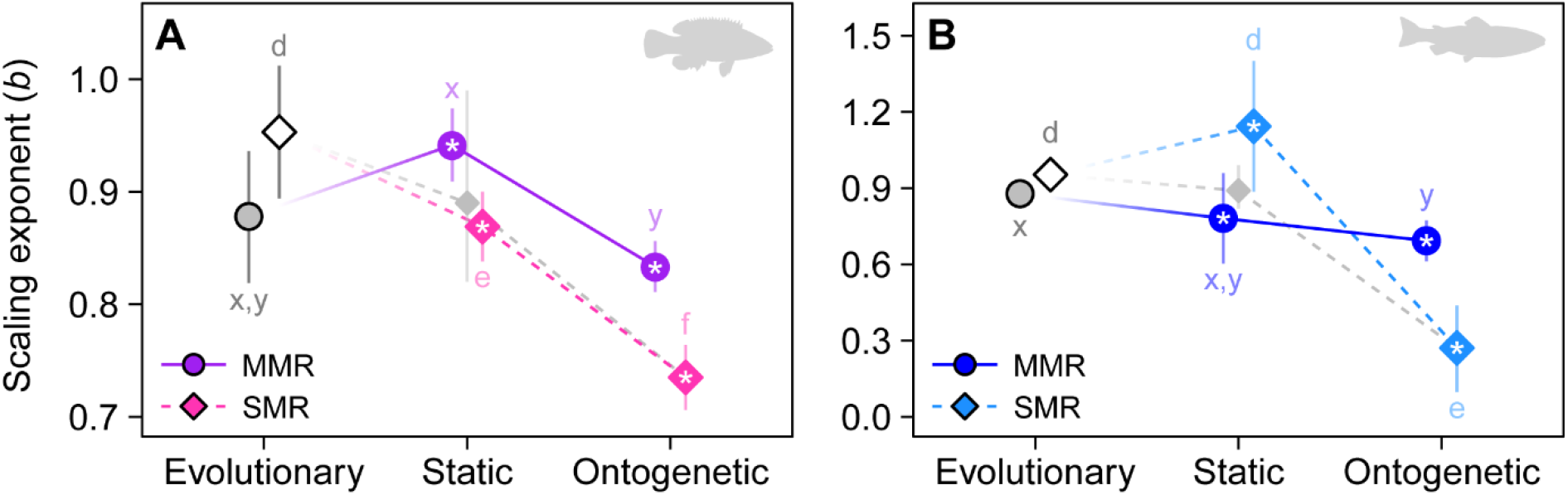
Comparison of metabolic rate scaling exponents between evolutionary, static, and ontogenetic levels. Exponents (*b*; ± 95% CI error bars) are for the scaling relationships shown in Fig. 1. Static and ontogenetic scaling exponents are shown for cunner (**A**) and brown trout (**B**) in coloured points. Different lower-case letters inside the panels indicate significant differences (p < 0.017) between exponents for either SMR or MMR; pairs of asterisks indicate significant differences between SMR and MMR within each taxonomic level (p < 0.05). The mean static *b* for SMR of 0.89 for 16 fishes (Jerde et al. 2019) is included for comparison (grey diamonds) and supports a general decrease in *b* for SMR from evolutionary across static to ontogenetic levels (grey dashed lines). Note the different y-axis scale on panel B (to accommodate the low ontogenetic *b* for trout).

There are no effects of either maximum body mass or relative body mass (i.e. at what body mass metabolic rate was measured, relative to the species’ maximum body mass) on the scaling exponent for either SMR or MMR (PGLS, interactions between measured mass and estimated maximum or relative mass; 0.664 ≥ p ≥ 0.171), or the scaling coefficient (intercept) for SMR or MMR (PGLS, effects of estimated maximum or relative mass; 0.104 ≥ p ≥ 0.095).

Scaling exponents for SMR of individual brown trout, which were fed a restricted ration of ~0.6% of their body mass daily, correlate positively with the fish’s growth rates (p < 0.0001; Fig. 3). There are no correlations between *b*_SMR_ and growth rate for cunner (which were fed an *ad libitum* ration of ~2.5% of their body mass daily), or for *b*_MMR_ and growth for either cunner or trout (Fig. 3).

**Figure 3.**
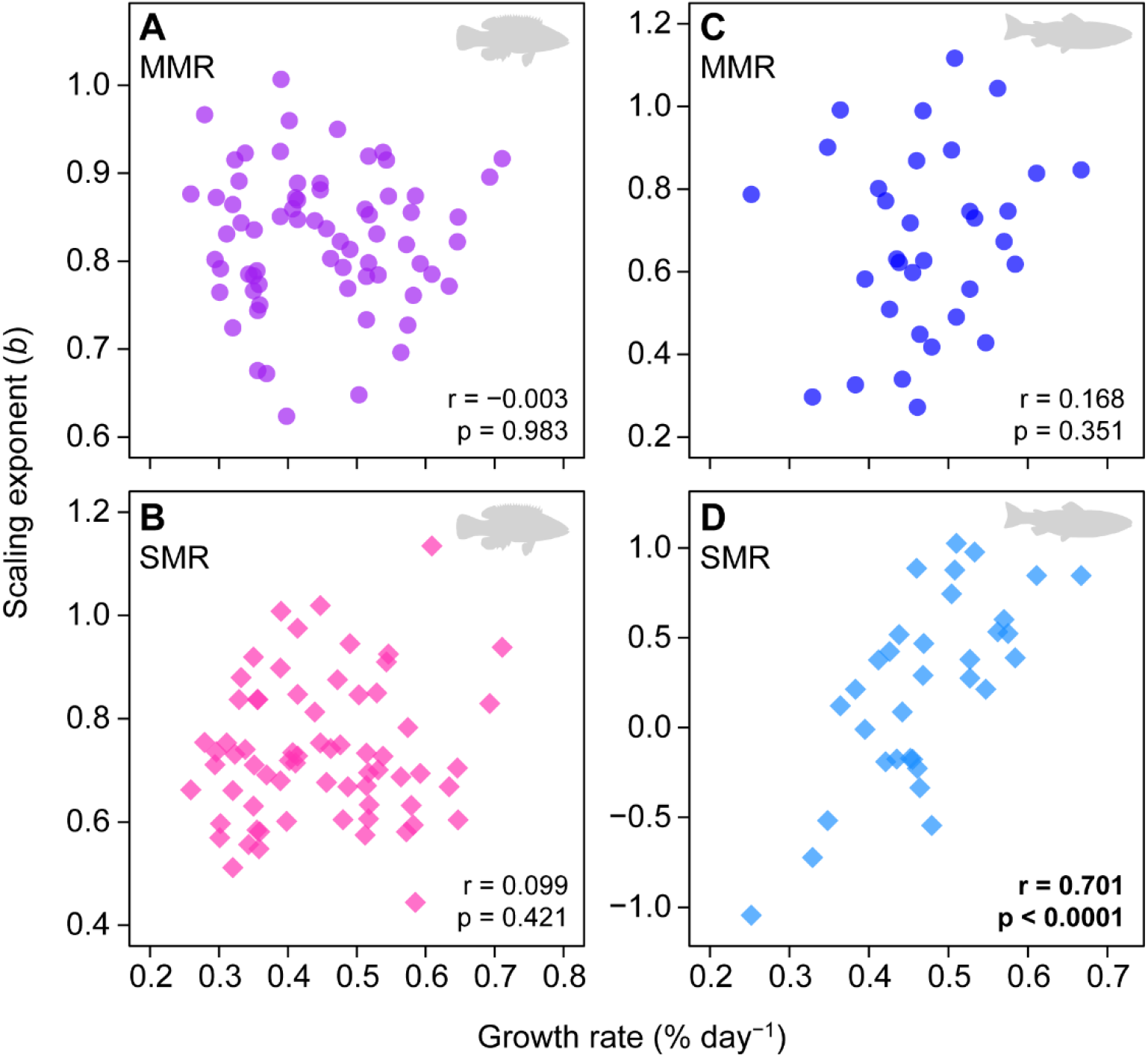
Relationships between metabolic rate scaling exponents and growth rates of individual fish. The observed variation in ontogenetic scaling exponents (*b*) for MMR (**A**, **C**) and SMR (**B**, **D**) of individual fish (cf. Fig. 1B-E) are unrelated to growth rates (A-C), except for SMR under restricted food intake (D). Cunner (A, B) were fed an *ad libitum* ration of ~2.5% of their body mass daily over the 10-month experiment (Norin & Gamperl 2018), while brown trout (C, D) were fed a restricted ration of ~0.6% of their body mass daily over the 15-week experiment (Norin & Malte 2011). Note that 11 out of 33 individuals in panel D exhibit negative scaling whereby SMR is reduced despite the fish growing; this is likely caused by the energetic constraint, which forced a reduction in SMR in some individuals (negative *b*) in order to maintain positive growth.

The scaling exponent for SMR within teleost orders increases significantly with the natural mortality rate of fish larvae of different species within those orders (p = 0.008; Fig. 4), while the exponent for MMR decreases (p = 0.028; Fig. 4).

**Figure 4.**
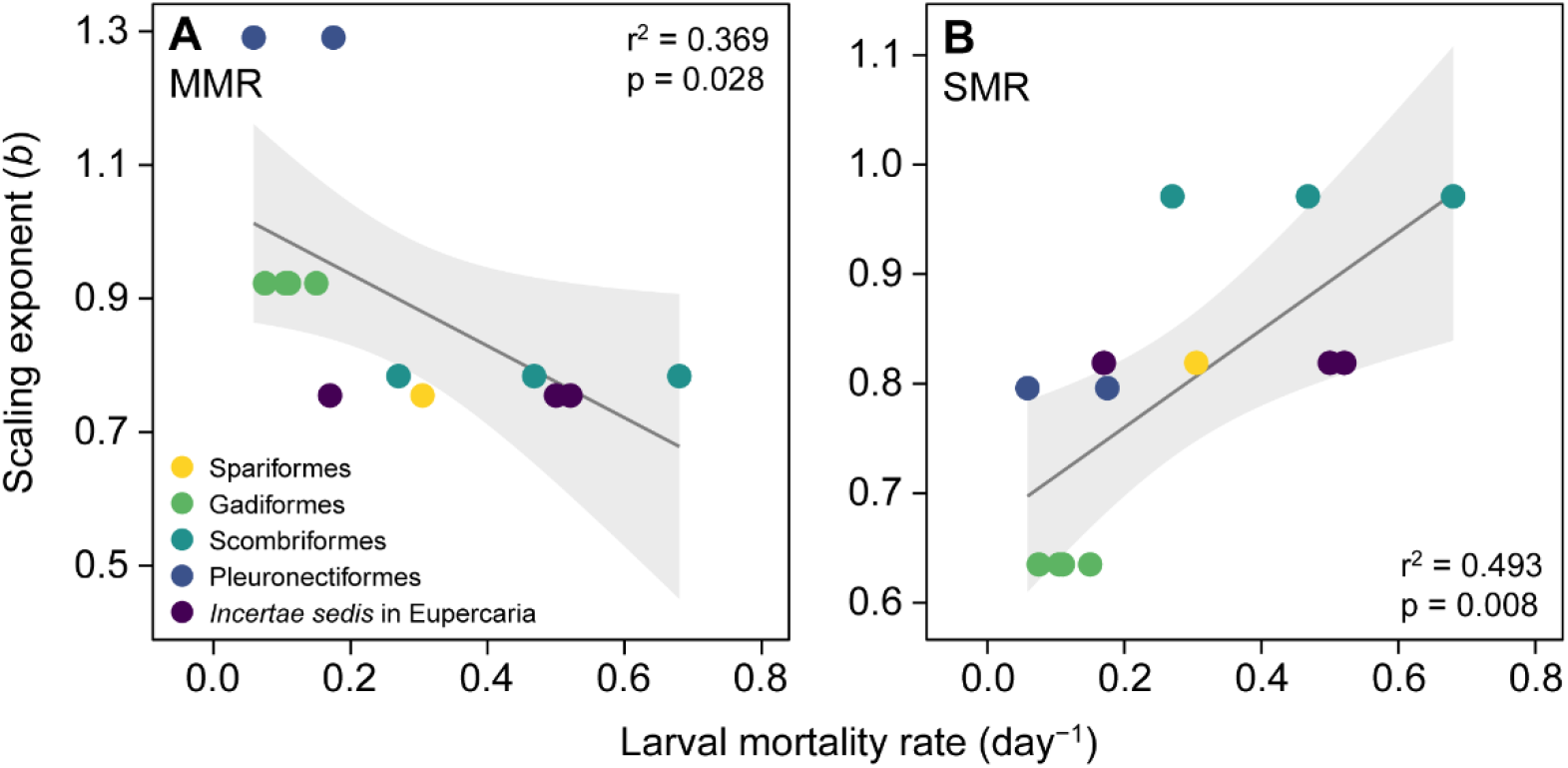
Metabolic rate scaling exponents vary predictably with natural mortality rates of fish larvae. Model-estimated scaling exponents for MMR (**A**) and SMR (**B**) of juvenile or adult fish from different teleost species (points) at the level of their order (colours) (see Fig. S1) change significantly with the natural larval mortality rates of the species, estimated in each species’ native marine environment.

## Discussion

I find that evolutionary (among-species) scaling of metabolic rate is steeper than ontogenetic (within-individual) scaling, in particular for standard (resting) metabolic rate, which scales with an exponent (*b*) near 1 among 134 species of fish (*b*_SMR_ = 0.953; Figs. 1 and 2). I also show that variation in ontogenetic scaling of SMR, but not MMR, can be explained by variation in growth rate when growth is restricted by limited food availability, with SMR scaling approximately isometrically (*b*_SMR_ ≈ 1) for the fastest growing individuals (Fig. 3). Finally, I find that *b*_SMR_ increases with natural mortality rates of fish larvae within orders for which data are available (Fig. 4) and suggest that selection on fast growth in early life is driving evolutionary scaling of SMR. While there is currently only reliable data on within-individual scaling available for two species of fish, and the broader generality of steeper evolutionary than ontogenetic scaling therefore needs further investigation, the finding of near-isometric evolutionary scaling is corroborated by a study on the scaling of routine metabolic rate across various groups of marine organisms (Kiørboe & Hirst 2014) and a recent study showing that maintenance metabolic rate scales with an exponent of 0.96 across eukaryotes (Hatton et al. 2019).

A somewhat surprising finding from re-analysing the brown trout data (Norin & Malte 2011) was a wide range of ontogenetic scaling exponents for SMR, with 11 out of 33 individuals exhibiting negative scaling whereby SMR was reduced despite the fish growing (Fig. 2E). This resulted in a mean ontogenetic *b*_SMR_ of only 0.271 within individuals. While such low and even negative scaling of metabolic rate is unusual, the strong correlation between *b*_SMR_ and growth rate of individual brown trout (Fig. 3D) indicate that the observed variation is real and biologically meaningful, and that negative scaling of metabolic rate (within individuals) is an overlooked phenomenon. A reduction in SMR in response to reduced food availability, as observed here for some trout, is a common observation among fish and other animals (Mueller & Diamond 2001; Fu et al. 2005; Wang et al. 2006; Van Leeuwen et al. 2012; Auer et al. 2016); however, this has previously been thought to increase the scaling exponent, as an inverse relationship between metabolic rate and its scaling exponent is predicted when energy availability is low (*sensu* the ‘metabolic-level boundaries hypothesis’; Glazier 2005, 2010). While this is found to be true at the static (among-individual) level where it has been studied in the past [reviewed by Glazier (2005)], and confirmed here (high static *b*_SMR_ of brown trout), my extended focus on within-individual scaling has revealed biological patterns that would have been overlooked with a more traditional species-level focus: metabolic scaling can be shallow and even negative at one level of biological organisation (within individuals) while being steep and positive at another (among individuals).

Much interest and effort has been devoted to metabolic growth models (von Bertalanffy 1957; Kooijman 2010; West et al. 2001; Ricklefs 2003; Hou et al. 2008, 2011; Barneche & Allen 2018; Marshall & White 2019), yet it remains debated if metabolic rate drives growth or if growth drives metabolic rate (Glazier et al. 2015). My findings indicate that the positive correlation between *b*_SMR_ and growth rate of brown trout is revealed by an energetic constraint, which forced a reduction in SMR in some individuals (negative *b*) in order to maintain positive growth; this, in turn, resulted in pronounced variation in metabolic scaling among individuals and, ultimately, a mean ontogentic *b*_SMR_ (0.271) much lower than the static and near-isometric *b*_SMR_ of 1.143 among the same individuals. Since the relationship between *b*_SMR_ and growth rate existed only for brown trout that were fed a restricted ration, compared to an *ad libitum* ration for cunner, this suggests that metabolic scaling is driven by growth rate – as also proposed and discussed by many others (Riisgård 1998; Glazier 2005, 2015, 2018; White et al. 2011; Tan et al. 2019) – but only when growth is constrained, as it usually is in nature where food is rarely readily available *ad libitum*.

Studies of individual variation in both physiology and behaviour are becoming more common in general (Roche et al. 2016), despite the additional work needed to phenotype the same animal multiple times, but there is still much to be explored. A specific prediction for future metabolic scaling research would be that differences between mean ontogenetic and static scaling exponents would increase with decreasing food availability. Since the cunner and brown trout examined here exhibited the same range of growth rates (~0.2–0.7% day^−1^), despite being fed very different rations, food availability arguably should be considered relative to the needs of each species examined when investigating the influence of growth on metabolic scaling at different levels of biological organisation.

Why then, is evolutionary scaling steeper than ontogenetic scaling, especially for SMR? Metabolic rates of fishes show a bi-phasic scaling pattern, with steeper and near-isometric scaling (*b* ≈ 1) in early life stages (Giguère et al. 1988; Post & Lee 1996; Killen et al. 2007). This is also the case for other animals with distinct larval and adult stages, such as copepods, insects, and marine invertebrates [reviewed by Glazier (2005)]. Since mortality is often very high in the early life stages of fishes (Houde 1997; Almany & Webster 2006; Hatton et al. 2019), it is likely that strong selection for fast growth out of the very vulnerable larval stage is driving the steep evolutionary scaling, pushing *b* towards 1, regardless of scaling patterns later in life. This is in agreement with early arguments that evolutionary brain size allometries arise from static or ontogenetic allometries when there is strong selection on body size (Gould 1975; Lande 1979; Riska & Atchely 1985). Steeper evolutionary than static scaling of brain size has been confirmed in mammals and birds and attributed to selection on early growth periods (Tsuboi et al. 2018), with brain size co-evolving with body size likely because genes affecting both brain and body size are expressed simultaneously (pleiotropy) early in life, whereas genes affecting only body size dominate later in life where selection for fast growth is relaxed (Riska & Atchley 1985). If the same applies to metabolic rate, this indicates that the steep evolutionary metabolic scaling observed here (Fig. 1A) is a by-product of selection for fast ontogenetic growth in the larval or early juvenile stages. If this potential pleiotropic gene effect on metabolic rate and body size is reduced later in life, this can also explain why metabolic rate appears to scale shallower at lower taxonomic levels (shallower average ontogenetic and static than evolutionary scaling; Fig. 2), and how there can be variation in *b* among species or taxa, as variable selection pressures and environmental effects acting on either metabolic rate or body size later in life can affect these traits independently of one another and cause variation in *b*.

While data on natural mortality of fish larvae are limited, the available evidence supports that strong selection (high natural mortality) in early life could indeed be driving *b*_SMR_ towards 1 (Fig. 4B). In other words, in taxa where mortality in early life is low, selection would not necessarily favour individuals with fast early-life growth, and those taxa would exhibit shallower metabolic scaling. That high mortality in early life may select for fast growth and steep scaling of metabolic rate has been proposed before (Epp & Lewis 1980; Glazier 2005), but with a focus on explaining static, species-level, scaling patterns. Glazier et al. (2011, 2020) also found evidence of shallower species-specific scaling of metabolic rate in freshwater amphipods naturally selected for slow growth; amphipods from springs where adult mortality is high due to size-selective fish predation exhibit lower *b*, relative to conspecifics from fish-less springs. These findings are in agreement with what I propose here, albeit in the opposite direction.

The idea of metabolic scaling patterns being driven by variation in growth also fits well with the patterns observed here, where the fastest growing juvenile brown trout, under a naturally-realistic (restricted) food regime, had scaling exponents for SMR around 1 (Fig. 3D), and faster growers would have higher survival in nature (Sogard 1997). The release from an energetic constraint in the *ad libitum* fed cunner arguably removed the relationship between ontogenetic metabolic scaling and growth rate in this species. Since growth of fishes is tightly related to SMR but not MMR (Auer et al. 2015, 2016; Norin et al. 2016), this could also explain why ontogenetic, static, and evolutionary scaling are more similar for MMR than for SMR (Fig. 2), and why *b*_SMR_ tends to be higher than *b*_MMR_ among species (i.e. the influence of growth rate on *b*_SMR_ does not affect *b*_MMR_). In turn, the reversal of this pattern at the lowest organisational level, evidenced by steeper ontogenetic scaling of MMR than SMR (Fig. 2), could be explained by the fitness advantage of keeping maintenance costs relatively low throughout an individual’s life (shallow ontogenetic scaling of SMR) while maximising scope for aerobic activity (steep ontogenetic scaling of MMR relative to SMR) (Killen et al. 2016). It is unclear why *b*_MMR_ appear to be negatively related to larval mortality rate (fig. 4), but it could be due to a negative relationships between *b*_SMR_ and *b*_MMR_ in early life, although no significant correlation between *b*_SMR_ and *b*_MMR_ is observed here at the order level (Pearson’s r = −0.290, p = 0.486; Fig. S2).

Given that the majority of fishes, along with most aquatic invertebrates, generally produce many (even millions) offspring that are spawned into the water or onto the substrate and left to fend for themselves, it is possible that the idea of steep evolutionary scaling being driven by selection for fast growth out of the vulnerable early-life stage is unique to aquatic ectotherms. Mortality rates in aquatic ectotherms scale steeper than for eukaryotes in general, with disproportionately high mortality rates of fish egg and larvae (Hatton et al. 2019). The idea is further supported by offspring size in the largest group of fishes, teleosts, being largely invariant across several orders of magnitude of adult asymptotic size (Neuheimer et al. 2015), indicating that all species would be subject to the same selective pressure in early life. On the other hand, it is also possible that selection on fast growth in early life is a universal driver of metabolic scaling among animals, but that the progressively shallower scaling of maintenance metabolic rate (standard metabolic rate in ectotherms, basal metabolic rate in endotherms) in reptiles, mammals, and birds, relative to fish and amphibians (White et al. 2006), is caused by a relaxation of this selection pressure and higher offspring survival in these lineages (e.g. due to parental care and relatively large offspring). A strong and positive genetic correlation between body mass and metabolic rate exists (White et al. 2019), and there is evidence that macroevolutionary relationships between metabolic rate and body mass result from multivariate selection on mass and metabolic rate (White et al. 2019). However, recent work indicates that there is no heritable genetic variation in the metabolic scaling exponent itself (Fossen et al. 2019; Beaman et al, preprint), whereas growth rate shows signs of heritable genetic variation (Beaman et al., preprint). Those findings also support the idea that selection on variation in body mass, because of variation in growth rate, could be an evolutionary driver of metabolic scaling.

Selection for fast growth in early life stages as a cause of steep evolutionary scaling could be explored in future work by comparing metabolic scaling of fish lineages that spawn many undefended eggs *vs*. those with viviparity (live-bearing) and/or parental care (e.g. mouth-brooders); species with viviparity or parental care produce relatively few offspring but these have higher survival and, presumably, relaxed selection on fast growth and thus lower scaling exponents, as also proposed by Glazier (2005). Glazier (2005, 2018) also speculated that evolutionary scaling of metabolic rate should be steeper in relatively large ectothermic species, because adult (post-maturational) growth could elevate metabolic rate relatively more in these species. I find no effect of either maximum body mass of fishes or their relative mass (i.e. at what body mass metabolic rate was measured, relative to the species’ maximum body mass) on metabolic rate and its mass-scaling, indicating that being a larger-growing species, or being measured at different post-larval stages of life, does not significantly influence metabolic scaling in fishes, whereas larval mortality rates do (Fig. 4).

The evolutionary scaling of MMR among the 114 fishes in the present study (*b*_MMR_ = 0.878) tends to be shallower than that of SMR (p = 0.077). The value of 0.878 for *b*_MMR_ is at the lower end of what has previously been reported among fewer (79) teleost species (*b*_MMR_ = 0.937, 95% CI = 0.864–1.011; Killen et al. 2016). While MMR is generally found to scale steeper than SMR at static (Brett 1965; Glazier 2014b) and ontogenetic (present study) levels, phylogenetically-informed work has reported similar evolutionary scaling exponents for SMR and MMR across both ectotherms (fishes, amphibians, and reptiles) and endotherms (mammals and birds) (Gillooly et al. 2017). The tendency observed here for shallower evolutionary scaling of MMR relative to SMR in fishes could indicate that the evolution of maximum body size in this group of animals is constrained by a decreasing aerobic scope (the difference between MMR and SMR) with increasing body mass. While it is unknown what the minimum aerobic scope required to successfully carry out life is, the metabolic rate required to sustain essential activities in the wild ranges from ~1.5 to 7 times maintenance metabolic rate across endo- and ectotherms (Peterson et al. 1990; Hammond & Diamond 1997), with a mode at 2-3 times maintenance metabolic rate (Deutsch et al. 2020). Thus, assuming that fish require an aerobic scope of at least twice their SMR (i.e. a factorial aerobic scope of two) to successfully carry out life (be physically active, digest, grow, and reproduce), this puts the maximum possible size of ray-finned fishes around the size of the largest extant teleost, the ocean sunfish (*Mola mola*), which has an estimated factorial aerobic scope of 1.9 at its maximum body mass of ~1,500 kg (based on the scaling relationships presented in Fig. 1). Although this could be coincidental, insufficient data are available on metabolic rates of large fishes to evaluate this, due to the logistical challenges of housing large fish in respirometry chambers. The on-going development of otolith-based proxies for retrospectively estimating lifetime metabolic rates and experienced environment (temperature and oxygen) of wild fishes (Chung et al. 2019, 2020; Limburg & Casini 2019) holds great potential for shedding light on the energetic requirements of fishes of all sizes.

In conclusion, my phylogenetically-informed analyses on metabolic rate data from 134 ray-finned fishes show that evolutionary scaling of especially SMR is near to isometric (*b*_SMR_ = 0.953), in agreement with previous phylogenetically-informed work on teleost fishes (*b*_SMR_ = 0.948; Killen et al. 2016) and eukaryotes in general (Kiørboe & Hirst 2014; Hatton et al. 2019), and that static and in particular ontogenetic (within-individual) scaling can be significantly shallower. The observed effects of growth (under restricted food availability) and natural mortality rates on metabolic scaling exponents support the idea that metabolic scaling is governed by ecological factors (Witting 1998; Killen et al. 2010; Harrison 2017; Uyeda et al. 2017; Glazier 2018; Hatton et al. 2019), rather than being set by physical or geometric constraints (West et al. 1997, 1999; Banavar et al. 2010) that, in turn, control ecological processes (Brown et al. 2004; Sibly et al. 2012).

## Author contributions

TN conceived the study; TN collected the data from the literature; TN analysed the data; TN wrote the paper and revised it.

## Data availability

Data and R script will be made available in the online supplementary material.

---

**Figure S1.**
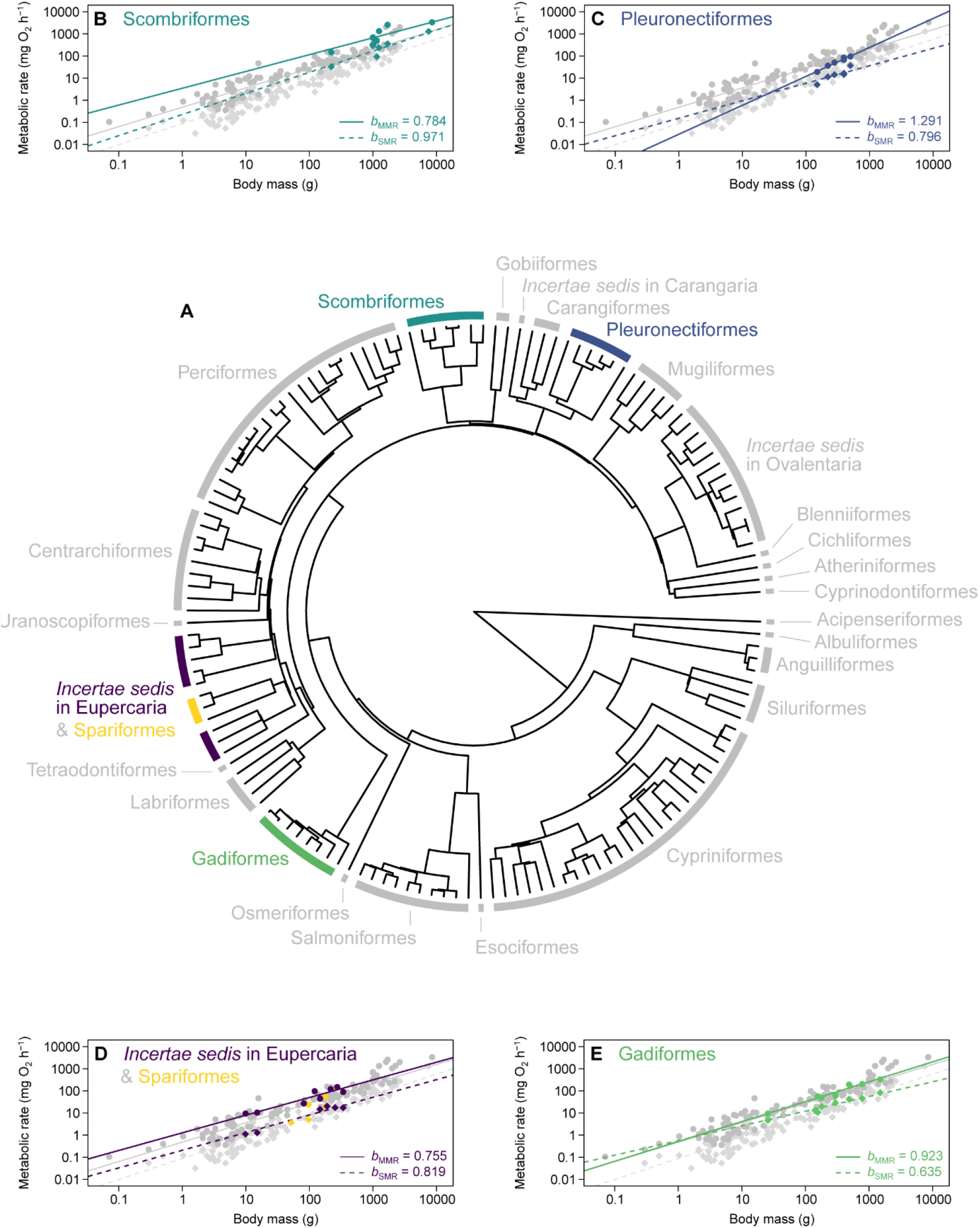
Phylogeny and metabolic rate scaling relationships for fishes from the present study. The phylogenetic tree (**A**) shows the evolutionary relationships between the 145 different species of ray-finned fish for which data for standard metabolic rate (SMR; 134 species) and maximum metabolic rate (MMR; 114 species) were used. The species are distributed among 27 orders, and the metabolic scaling relationships for species within five of these orders (**B-E**) are highlighted and colour coded for comparison with Fig. 4 in the main manuscript (note that Spariformes have been combined with *Incertae sedis* in Eupercaria due to their close taxonomic affiliation). Scaling exponents (*b*) for SMR and MMR of the species within each highlighted order are indicated on the panels, and fitted regression lines have been extended to the edges of the plots for easier visual comparison with the overall scaling relationships across all species (grey points and lines). Diamonds and dashed lines represent SMR while circles and solid lines represent MMR.

**Figure S2.**
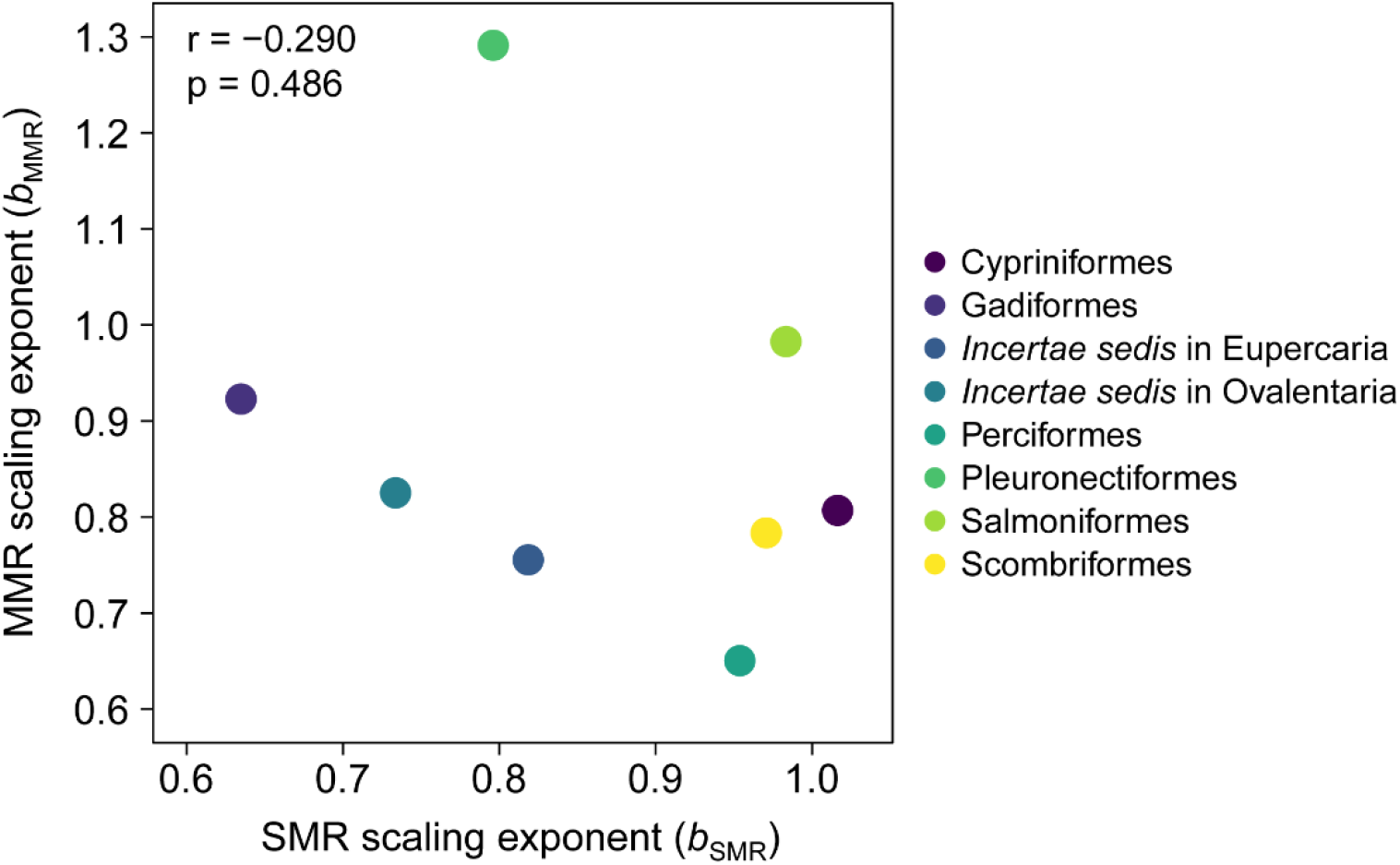
Correlation between scaling exponents for standard- and maximum metabolic rates of different teleost orders. Scaling exponents for both standard metabolic rate (*b*_SMR_) and maximum metabolic rate (*b*_MMR_) are model-derived estimates for orders with at least six species in the dataset. Note that the colour coding used here does not correspond to those used in Fig. S1 and Fig. 4 in the main manuscript, as only orders for which natural larval mortality rates were available in the literature are shown in those figures.

## References

Almany GR & Webster MS (2006) The predation gauntlet: early post-settlement mortality in reef fishes. Coral Reefs 25, 19–22.

Andersen KH (2019) Fish ecology, evolution, and exploitation: a new theoretical synthesis. Princeton University Press, Princeton, New Jersey.

Auer SK, Salin K, Rudolf AM, Anderson GJ & Metcalfe NB (2015) Flexibility in metabolic rate confers a growth advantage under changing food availability. J. Anim. Ecol. 84, 1405–1411.

Auer SK, Salin K, Rudolf AM, Anderson GJ & Metcalfe NB (2016) Differential effects of food availability on minimum and maximum rates of metabolism. Biol. Lett. 12, 20160586.

Banavar JR, Moses ME, Brown JH, Damuth J, Rinaldo A, Sibly RM & Maritan A (2010) A general basis for quarter-power scaling in animals. Proc. Natl. Acad. Sci. USA 107, 15816–15820.

Barneche DR & Allen AP (2018) The energetics of fish growth and how it constrains food-web trophic structure. Ecol. Lett. 21, 836–844.

Bates D, Maechler M, Bolker B & Walker S (2015) Fitting linear mixed-effects models using lme4. J. Stat. Softw. 67, 1–48.

Beaman JE, Ortiz-Barrientos D, Monro K, Hall MD & White CR (preprint) Metabolic scaling has diversified among species, despite an evolutionary constraint within species. bioRxiv 2020.05.26, 117846, doi:https://doi.org/10.1101/2020.05.26.117846.

Brett JR (1965) The relation of size to rate of oxygen consumption and sustained swimming speed of sockeye salmon (Oncorhynchus nerka). J. Fish. Res. Bd. Canada 22, 1491–1501.

Brody S (1945) Bioenergetics and growth. Reinhold, New York.

Brown JH, Gillooly JF, Allen AP, Savage VM & West GB (2004) Toward a metabolic theory of ecology. Ecology 85, 1771–1789.

Brown JH, Hall CAS & Sibly RM (2018) Equal fitness paradigm explained by a trade-off between generation time and energy production rate. Nat. Ecol. Evol. 2, 262–268

Chang J, Rabosky DL, Smith SA, Alfaro ME (2019) An R package and online resource for macroevolutionary studies using the ray-finned fish tree of life. Methods Ecol. Evol. 10, 1118–1124.

Cheung WWL, Sarmiento JL, Dunne J, Frölicher TL, Lam, Palomares, Watson R & Pauly D (2013) Shrinking of fishes exacerbates impacts of global ocean changes on marine ecosystems. Nat. Clim. Change 3, 254–258.

Cheverud JM (1982) Relationships among ontogenetic, static, and evolutionary allometry. Am. J. Phys. Antropol. 59, 139–149.

Chung M-T, Trueman CN, Godiksen JA, Holmstrup ME & Grønkjær P (2019) Field metabolic rates of teleost fishes are recorded in otolith carbonate. Commun. Biol. 2:24

Chung M-T, Jørgensen K-EM, Trueman CN, Knutsen H, Jorde PE & Grønkjær P (2020) First measurements of field metabolic rate in wild juvenile fishes show strong thermal sensitivity but variations between sympatric ecotypes. Oikos, doi: 10.1111/oik.07647

Clarke A & Johnston NM (1999) Scaling of metabolic rate with body mass and temperature in teleost fish. J. Anim. Ecol. 68, 893–905.

Davis TLO, Lyne V & Jenkins GP (1991) Advection, dispersion and mortality of a patch of southern Bluefin tuna larvae Thunnus maccoyii in the East Indian Ocean. Mar. Ecol. Prog. Ser. 73, 33–45.

DeLong JP, Okie JG, Moses ME, Sibly RM & Brown JH (2010) Shifts in metabolic scaling, production, and efficiency across major evolutionary transitions of life. Proc. Natl. Acad. Sci. USA 107, 12941–12945.

Deutsch C, Ferrel A, Seibel B, Pörtner H-O & Huey RB (2015) Climate change tightens a metabolic constraint on marine habitats. Science 348, 1132–1135.

Deutsch C, Pell JL & Seibel B (2020) Metabolic trait diversity shapes marine biogeography. Nature 585, 557–562.

Epp RW & Lewis WM Jr (1980) The nature and ecological significance of metabolic changes during the life history of copepods. Ecology 61, 259–264.

Fossen EIF, Pélabon C & Einum S (2019) Genetic and environmental effects on the scaling of metabolic rate with body size. J. Exp. Biol. 222, jeb193243.

Freckleton RP, Harvey PH & Pagel M (2002) Phylogenetic analysis and comparative data: a test and review of evidence. Am. Nat. 160, 712–726.

Froese R & Pauly D (2019) FishBase. World Wide Web electronic publication. www.fishbase.org, version 12/2019.

Fu SJ, Xie XJ & Cao ZD (2005) Effect of fasting on resting metabolic rate and postprandial metabolic response in *Silurus meridionalis*. J. Fish Biol. 67, 279–285.

Giguère LA, Côté B & St-Pierre J-F (1988) Metabolic rates scale isometrically in larval fishes. Mar. Ecol. Prog. Ser. 50, 13–19.

Gillooly JF, Gomez JP & Mavrodiev EV (2017) A broad-scale comparison of aerobic activity levels in vertebrates: endotherms versus ectotherms. Proc. R. Soc. B 284, 20162328.

Glazier DS (2005) Beyond the ‘3/4-power law’: variation in the intra- and interspecific scaling of metabolic rate in animals. Biol. Rev. 80, 611–662.

Glazier DS (2010) A unifying explanation for diverse metabolic scaling in animals and plants. Biol. Rev. 85, 111–138.

Glazier DS (2014a) Metabolic scaling in complex living systems. Systems 2, 451–540.

Glazier DS (2014b) Scaling of metabolic scaling within physical limits. Systems 2, 425–450.

Glazier DS (2015) Is metabolic rate a universal ‘pacemaker’ for biological processes? Biol. Rev. 90, 377–407.

Glazier DS (2018) Rediscovering and reviving old observations and explanations of metabolic scaling in living systems. Systems 6, systems6010004.

Glazier DS, Butler EM, Lombardi SA, Deptola TJ, Reese AJ & Satterthwaite EV (2011) Ecological effects on metabolic scaling: amphipod responses to fish predators in freshwater springs. Ecol. Monogr. 81, 599–618.

Glazier DS, Borrelli JJ & Hoffman CL (2020) Effects of fish predators on the mass-related energetics of a keystone freshwater crustacean. Biology 9, biology9030040.

Gould SJ (1966) Allometry and size in ontogeny and phylogeny. Biol. Rev. 41, 587–640.

Gould SJ (1975) Allometry in primates, with emphasis on scaling and the evolution of the brain. In Approaches to primate paleobiology (Szalay F, ed.). Karger, Basel, pp. 244–292.

Hammond KA & Diamond J (1997) Maximal sustained energy budgets in humans and animals. Nature 386, 457–462.

Harrison JF (2017) Do performance-safety tradeoffs cause hypometric metabolic scaling in animals? Trends Ecol. Evol. 32, 653–664.

Hatton IA, Dobson AP, Galbraith ED & Loreau M (2019) Linking scaling laws across eukaryotes. Proc. Natl. Acad. Sci. USA 116, 21616–21622.

Hou C, Zuo W, Moses ME, Woodruff WH, Brown JH & West GB (2008) Energy uptake and allocation during ontogeny. Science 322, 736–739.

Hou C, Bolt KM & Bergman A (2011) A general model for ontogenetic growth under food restriction. Proc. R. Soc. B 278, 2881–2890.

Houde ED (1997) Patterns and trends in larval-stage growth and mortality of teleost fish. J. Fish Biol. 51, 52–83.

Houde ED & Zastrow CE (1993) Ecosystem- and taxon-specific dynamic and energetics properties of larval fish assemblages. Bull. Mar. Sci. 53, 290–335.

Jerde CL, Kraskura K, Elison EJ, Csik SR, Stier A & Taper ML (2019) Strong evidence for an intraspecific metabolic scaling coefficient near 0.89 in fish. Front. Physiol. 10, 1166.

Killen SS, Costa I, Brown JA & Gamperl AK (2007) Little left in the tank: metabolic scaling in marine teleosts and its implications for aerobic scope. Proc. R. Soc. B 274, 431–438.

Killen SS, Atkinson D & Glazier DS (2010) The intraspecific scaling of metabolic rate with body mass in fishes depends on lifestyle and temperature. Ecol. Lett. 13, 184–193.

Killen SS, Glazier DS, Rezende EL, Clark TD, Atkinson D, Willener AST & Halsey LG (2016) Ecological influences and morphological correlates of resting and maximal metabolic rates across teleost fish species. Am. Nat. 187, 592–606.

Killen SS, Norin T & Halsey LG (2017) Do method and species lifestyle affect measures of maximum metabolic rate in fishes? J. Fish Biol. 90, 1037–1046.

Kiørboe T & Hirst AG (2014) Shifts in mass scaling of respiration, feeding, and growth rates across life-form transitions in marine pelagic organisms. Am. Nat. 183, E118–E130.

Kleiber M (1932) Body size and metabolism. Hilgardia 6, 315–353.

Kleiber M (1947) Body size and metabolic rate. Physiol. Rev. 27, 511–541.

Kolokotrones T, Savage V, Deeds EJ & Fontana W (2010) Curvature in metabolic scaling. Nature 464, 753–756.

Kooijman SALM (2010) Dynamic energy budget theory for metabolic organisation (3rd ed.). Cambridge University Press, Cambridge.

Kozłowski J & Weiner J (1997) Interspecific allometries are by-products of body size optimization. Am. Nat. 149, 352–380.

Kozłowski J, Konarzewski M & Gawelczyk AT (2003) Cell size as a link between noncoding DNA and metabolic rate scaling. Proc. Natl. Acad. Sci. USA 100, 14080–14085.

Kuznetsova A, Brockhoff PB & Christensen RHB (2017) lmerTest package: tests in linear mixed effects models. J. Stat. Softw. 82, 1–26.

Lande R (1979) Quantitative genetic analysis of multivariate evolution, applied to brain–body size allometry. Evolution 33, 402–416.

Lande R (1985) Genetic and evolutionary aspects of allometry. In Size and scaling in primate biology (Jungers WL, ed.). Plenum, New York, pp. 21–32.

Limburg KE & Casini M (2019) Otolith chemistry indicates recent worsened Baltic cod condition is linked to hypoxia exposure. Biol. Lett. 15, 20190352.

Marshall DJ & White CR (2019) Have we outgrown the existing models of growth? Trends Ecol. Evol. 34, 102–111.

McGurk MD (1986) Natural mortality of marine pelagic fish eggs and larvae: role of spatial patchiness. Mar. Ecol. Prog. Ser. 34, 227–242.

Mueller P & Diamond J (2001) Metabolic rate and environmental productivity: well-provisioned animals evolved to run and idle fast. Proc. Natl. Acad. Sci. USA 98, 12550–12554.

Neuheimer AB, Hartvig M, Heuschele J, Hylander S, Kiørboe T, Olsson KH, Sainmont J & Andersen KH (2015) Adult and offspring size in the ocean over 17 orders of magnitude follows two life history strategies. Ecology 96, 3303–3311.

Norin T & Malte H (2011) Repeatability of standard metabolic rate, active metabolic rate and aerobic scope in young brown trout during a period of moderate food availability. J. Exp. Biol. 214, 1668–1675.

Norin T, Malte H & Clark TD (2016) Differential plasticity of metabolic rate phenotypes in a tropical fish facing environmental change. Funct. Ecol. 30, 369–378.

Norin T & Gamperl AK (2018) Metabolic scaling of individuals vs. populations: evidence for variation in scaling exponents at different hierarchical levels. Funct. Ecol. 32, 379–388.

Norin T & Speers-Roesch B (2020) Chapter 10 – Metabolism. In The Physiology of Fishes, 5^th^ed. (Currie S & Evans DH, eds.). CRC Press, Taylor & Francis Group, Boca Raton, FL, pp. 128–141.

Paradis A & Schliep K (2019) ape 5.0: an environment for modern phylogenetics and evolutionary analyses in R. Bioinformatics 35, 526–528.

Pélabon C, Bolstad GH, Egset CK, Cheverud JM, Pavlicev M & Rosenqvist G (2013) On the relationship between ontogenetic and static allometry. Am. Nat. 181, 195–212.

Pepin P (1991) Effect of temperature and size on development, mortality, and survival rates of the pelagic early life history stages of marine fish. Can. J. Fish. Aquat. Sci. 48, 503–518.

Peterson CC, Nagy KA & Diamond J (1990) Sustained metabolic scope. Proc. Natl. Acad. Sci USA 87, 2324–2328.

Post JR & Lee JA (1996) Metabolic ontogeny of teleost fishes. Can. J. Fish. Aquat. Sci. 53, 910–923.

R Core Team (2020) R: a language and environment for statistical computing. R Foundation for Statistical Computing, Vienna, Austria. URL: https://www.R-project.org/.

Rabosky DL, Chang J, Title PO, Cowman PF, Sallan L, Friedman M, Kaschner K, Garilao C, Near TJ, Coll M & Alfaro ME (2018) An inverse latitudinal gradient in speciation rate for marine fishes. Nature 559, 392–395.

Ricklefs RE (2003) Is rate of ontogenetic growth constrained by resource supply or tissue growth potential? A comment on West et al.’s model. Funct. Ecol. 17, 384–393.

Riisgård HU (1998) No foundation of a “3/4 power scaling law” for respiration in biology. Ecol. Lett. 1, 71–73.

Riska B & Atchley WR (1985) Genetics of growth predict patterns of brain-size evolution. Science 229, 668–671.

Roche DG, Careau V & Binning SA (2016) Demystifying animal ‘personality’ (or not): why individual variation matters to experimental biologists. J. Exp. Biol. 219, 3832–3843.

Rubner M (1883) Über den einfluss der körpergrösse auf stoff-und kraftwechsel. Zeitschr. f. Biol. 19, 535–562.

Savage VM, Gillooly JF, Woodruff WH, West GB, Allen AP, Enquist BJ & Brown JH (2004) The predominance of quarter-power scaling in biology. Funct. Ecol. 18, 257–282.

Schmidt-Nielsen K (1984) Scaling: why is animal size so important? Cambridge University Press, Cambridge.

Sibly RM, Brown JH & Kodric-Brown A (2012) Metabolic ecology: a scaling approach. Wiley-Blackwell, John Wiley & Sons Ltd, West Sussex.

Sibly RM & Brown JH (2020) Toward a physiological explanation of juvenile growth curves. J. Zool. 311, 286–290.

Sogard SM (1997) Size-selective mortality in the juvenile stage of teleost fishes: a review. Bull. Mar. Sci. 60, 1129–1157.

Tan H, Hirst AG, Glazier DS & Atkinson D (2019) Ecological pressures and the contrasting scaling of metabolism and body shape in coexisting taxa: cephalopods versus teleost fish. Phil. Trans. R. Soc. B 374, 20180543.

Tsuboi M, ven der Bijl, Kopperud BT, Erritzøe J, Voje KL, Kotrschal A, Yopak KE, Collin SP, Iwaniuk AN & Kolm N (2018) Breakdown of brain–body allometry and the encephalization of birds and mammals. Nat. Ecol. Evol. 2, 1492–1500.

Uyeda JC, Pennell MW, Miller ET, Maia R & McClain CR (2017) The evolution of energetic scaling across the vertebrate tree of life. Am. Nat. 190, 185–199.

Voje KL, Hansen TF, Egset CK, Bolstad GH & Pélabon C (2013) Allometric constraints and the evolution of allometry. Evolution 68, 866–885.

van de Pol M & Wright J (2009) A simple method for distinguishing within-versus between-subject effects using mixed models. Anim. Behav. 77, 753–758.

Van Leeuwen TE, Rosenfeld JS & Richards JG (2012) Effects of food ration on SMR: influence of food consumption on individual variation in metabolic rate in juvenile coho salmon (*Oncorhynchus kisutch*). J. Anim. Ecol. 81, 395–402.

von Bertalanffy L (1957) Quantitative laws in metabolism and growth. Q. Rev. Biol. 32, 217–231.

Wang T, Hung CCY & Randall DJ (2006) The comparative physiology of food deprivation: from feast to famine. Annu. Rev. Physiol. 68, 223–251.

West GB, Brown JH & Enquist BJ (1997) A general model for the origin of allometric scaling laws in biology. Science 276, 122–126.

West GB, Brown JH & Enquist BJ (1999) The fourth dimension of life: fractal geometry and allometric scaling of organisms. Science 284, 1677–1679.

West GB, Brown JH & Enquist BJ (2001) A general model for ontogenetic growth. Nature 413, 628–631.

White CR & Seymour RS (2003) Mammalian basal metabolic rate is proportional to body mass^2/3^. Proc. Natl. Acad. Sci. USA 100, 4046–4049.

White CR, Phillips NF & Seymour RS (2006) The scaling and temperature dependence of vertebrate metabolism. Biol. Lett. 2, 125–127.

White CR, Cassey P & Blackburn TM (2007) Allometric exponents do not support a universal metabolic allometry. Ecology 88, 315–323.

White CR, Kearney MR, Matthews PGD, Kooijman SALM & Marshall DJ (2011) A manipulative test of competing theories for metabolic scaling. Am. Nat. 178, 746–754.

White CR, Marshall DJ, Alton LA, Arnold PA, Beaman JE, Bywater CL, Condon C, Crispin TS, Janetzki A, Pirtle E, Winwood-Smith HS, Angilletta MJ Jr, Chenoweth SF, Franklin CE, Halsey LG, Kearney MR, Portugal SJ & Ortiz-Barrientos D (2019) The origin and maintenance of metabolic allometry in animals. Nat. Ecol. Evol. 3, 598–603.

Witting L (1998) Body mass allometries caused by physiological or ecological constraints? Trends Ecol. Evol. 13, 25.

